# An Antibiotic-Treated Mouse Model Reveals the Progression of Intestinal Botulism

**DOI:** 10.64898/2026.06.17.732786

**Authors:** Takuhiro Matsumura, Nobuhide Kobayashi, Sho Amatsu, Kazuki Saito, Aki Yamaguchi, Kanon Iwata, Yukako Fujinaga

**Author notes:** Adress correspondence to Takuhiro Matsumura;, or Yukako Fujinaga.

## Abstract

Intestinal botulism, including infant botulism and adult intestinal toxemia botulism, is a life-threatening disease caused by intestinal infection with *Clostridium botulinum* (*Cb*) spores. Infants are particularly susceptible to *Cb* infection because of their immature gut microbiota, whereas healthy adults are generally protected by gut microbiota-mediated colonization resistance. Thus, immature gut microbiota or gut dysbiosis is thought to permit *Cb* colonization. However, the mechanisms underlying colonization resistance and disease progression remain poorly characterized. Here, using adult mice with antibiotic-induced dysbiosis, we established a model for studying *Cb* infection and characterized intestinal colonization, bacterial expansion, botulinum neurotoxin (BoNT) accumulation, and disease progression during intestinal botulism. Intestinal botulism developed in antibiotic-treated mice after intragastric administration of strain 62A spores, whereas untreated adult mice showed no symptoms. In antibiotic-treated mice, *Cb* expanded over time in fecal samples, followed by accumulation of serotype A BoNT (BoNT/A), which correlated with the progression of botulism symptoms. *Cb* growth and BoNT/A accumulation occurred mainly in the cecum and colon, but not in the small intestine. Furthermore, this mouse model waapplicable to the analysis of intestinal botulism caused by other *Cb* strains, including 7I03-H, Okra, and Osaka05. Taken together, our findings clarify key aspects of intestinal botulism pathogenesis. This mouse model may provide a useful platform for elucidating the mechanisms underlying intestinal botulism, including infant botulism, and for developing novel therapeutic strategies.

**Importance:** Botulism is a life-threatening disease caused by botulinum neurotoxin, one of the most powerful toxins known. In intestinal botulism, Clostridium botulinum spores enter the intestine, grow there, and release toxin inside the body. However, the detailed process by which intestinal botulism develops remains poorly understood. In this study, we established a mouse model that allows the progression of intestinal botulism to be examined over time. Using this model, we showed that bacterial growth in the intestine, accumulation of biologically active toxin, and worsening of symptoms are closely linked. We also identified the cecum and colon as the main sites of bacterial colonization and toxin accumulation. This model can be used to analyze multiple C. botulinum strains with different characteristics and provides a useful tool for understanding how intestinal botulism develops and for guiding future preventive and therapeutic strategies.

## Introduction

Botulism is a severe neuroparalytic disease caused by botulinum neurotoxin (BoNT), which is produced by *Clostridium botulinum* (*Cb*) and related species (1,2), and can be fatal in approximately 40–50% of untreated cases (3). BoNT cleaves SNAREs at neuromuscular junction, thereby inhibiting neurotransmitter release and causing a paralysis (1,2). BoNTs have classically been classified into seven serotypes, A–G, with multiple subtypes (4,5) and types A, B, E, and F cause human botulism (2). Intestinal botulism, including infant botulism and adult intestinal toxemia botulism, together with foodborne botulism, account for the majority of botulism cases. Although adult intestinal toxemia botulism is exceedingly rare, more than 100 cases of infant botulism are reported annually in the United States (6–8). Foodborne botulism also remains an important public health concern, with outbreaks continuing to be reported in Southeast Asia (9–12). Foodborne botulism is caused by ingestion of BoNT contaminated food, whereas intestinal botulism is caused by ingestion of *Cb* spores. In intestinal botulism, ingested spores germinate in the intestine, where the vegetative cells grow and produce BoNT, leading to flaccid paralysis. Normal adult mice are resistant to intragastric injection of botulinum spores, whereas germ-free adult mice and neonatal mice are susceptible to infection, suggesting that mature intestinal microbiota provides colonization resistance against botulinum spores (13–16). Furthermore, adult mice orally treated with antibiotics are susceptible to botulinum spores, indicating that antibiotic-induced dysbiosis disrupts intestinal colonization resistance against *Cb* (17,18).

Consistent with these experimental findings, adult intestinal toxemia botulism has been reported in patients receiving antimicrobial therapy. By 2020, more than 20 cases had been described in the literature, and associations with a variety of antibiotics have been suggested (19–27). As described above, previous studies established the importance of intestinal microbiota in resistance to *Cb* infection (13–18). However, the detailed pathological processes following intestinal colonization, including the temporal relationship among *Cb* growth, toxin production, anatomical sites of toxin production, and clinical disease progression, remain poorly characterized.

Here, we established a mouse model to investigate the pathogenesis of intestinal botulism in detail. We analyzed disease progression during infection by quantifying BoNT levels using sandwich ELISA and mouse bioassay, measuring bacterial abundance by colony-forming unit (CFU) assay and RT-qPCR, and evaluating clinical symptoms over time. Using this system, we demonstrated that the abundance of *Cb* and BoNT in fecal samples correlated with disease progression and identified cecum and colon as the primary sites of *Cb* proliferation and toxin production. Furthermore, we demonstrated that this system is broadly applicable to Group I *Cb* strains. Collectively, by establishing a quantitative and reproducible mouse model, we clarified how intestinal *Cb* colonization, BoNT accumulation, and disease progression are linked during intestinal botulism.

## Results

### Antibiotic-induced intestinal dysbiosis contributes to the onset of intestinal botulism

Various antibiotics can induce dysbiosis but alter the intestinal microbiota in distinct ways (28). Consequently, different antibiotic-induced microbiota compositions may influence susceptibility to bacterial infections (29). Sugiyama and colleagues have demonstrated that erythromycin/kanamycin (EK)- or metronidazole (Met)-treated mice become susceptible to *Cb* infection (17,18). We first examined whether pretreatment with different antibiotics before intragastric administration of strain 62A spores (1 × 10^6^ CFU) could induce botulism in mice. For this purpose, we tested antibiotic regimens previously used in intestinal botulism models, including EK and Met, as well as a newly selected regimen consisting of ampicillin and vancomycin (AV). Mice treated with EK or with Met did not show clear symptoms of botulism, with disease score of 0 or less than 1. In contrast, mice treated with AV developed botulism and died within 7 days (Fig. S1a). Based on these results, we selected the AV for subsequent experiments (Fig. 1a). We next confirmed whether AV treatment induced intestinal dysbiosis. Quantitative PCR analysis showed that the total bacterial abundance in the intestine was significantly reduced in AV-treated mice compared with control mice at day 0, indicating that AV treatment disrupted the intestinal microbiota (Fig. 1b). After intragastric administration of strain 62A spores, AV-treated mice showed clear botulism symptoms on day 5 and died within 8 days, whereas control mice showed no symptoms throughout the observation period (Fig. 1c,d). Under these conditions, *Cb* colonized all AV-treated mice, and all colonized mice developed botulism that was ultimately fatal. Although administration of 1 × 10 □CFU of spores also caused infection and disease development (Fig. S1b), 1 × 10 □ CFU resulted in more consistent infection and disease progression. These results indicate that AV-induced dysbiosis provides a susceptible intestinal environment that permits *Cb* colonization and subsequent development of intestinal botulism.

**Figure 1.**
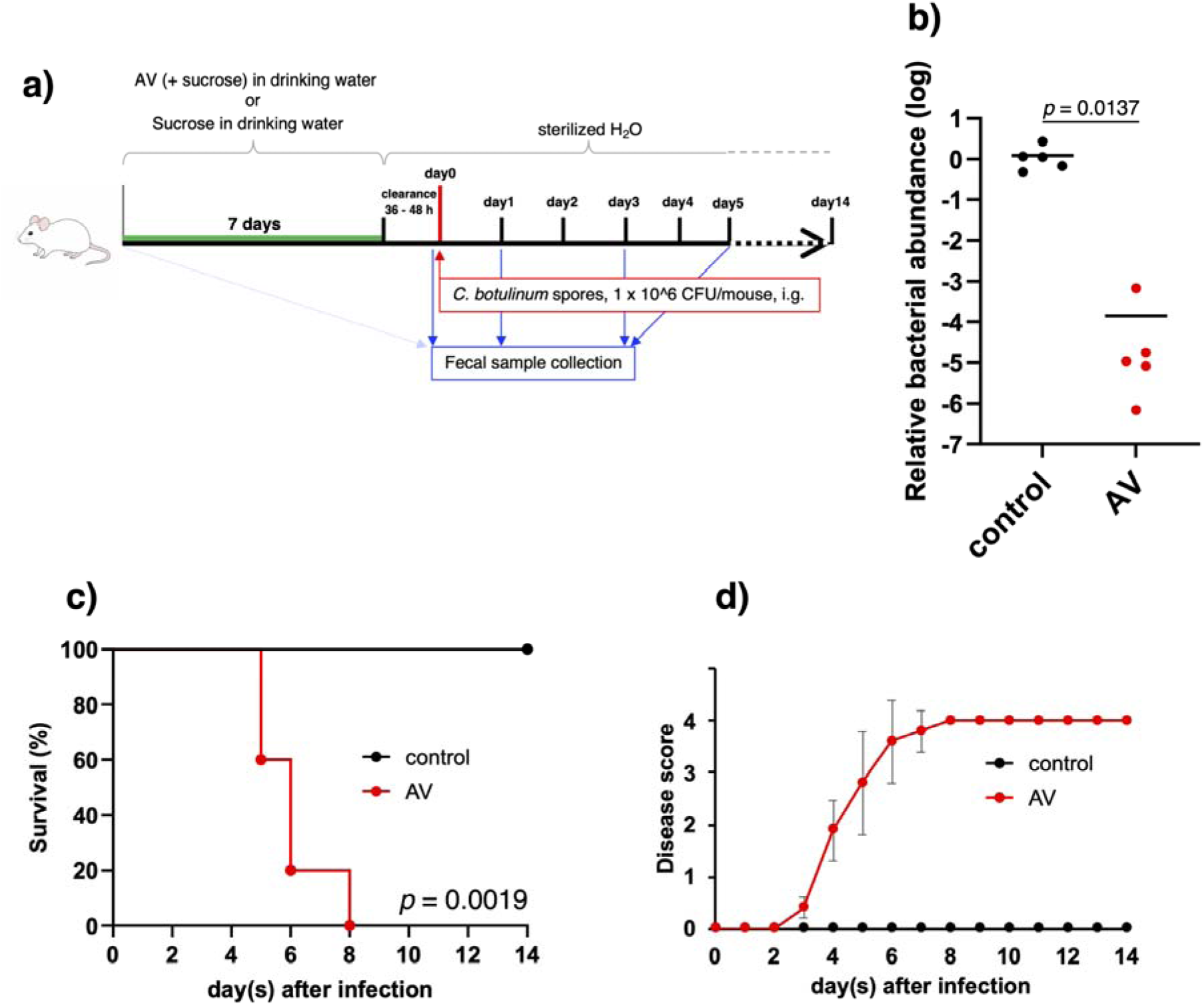
Intestinal botulism using antibiotic-treated mice. **a** Experimental design of the intestinal botulism mouse model. Antibiotic solution containing ampicillin and vancomycin (AV) or control sucrose solution was administered to BALB/c adult mice (*n* = 5) via drinking water for 1 week. The drinking water was then replaced with sterile water for 36–48 h before spore administration to allow antibiotic washout. Spore suspensions containing 1 × 10 CFU were intragastrically administered to AV-treated or control mice. **b** Total bacterial abundance in fecal samples on day 0 was analyzed by qPCR using primers targeting the bacterial 16S rRNA gene. Each dot represents an individual mouse. The data were analyzed by two-tailed Welch’s t-test. **cd** Spore-administered mice (*n* = 5) were monitored for survival (**c**) and disease scores (**d**) during a 2-week observation period. **c** Statistical analysis was performed using the log-rank test. **d** Data are presented as mean ± s.d. Data are representative of two independent experiments.

### *Cb* proliferates in the intestine of AV-treated mice

To analyze *Cb* growth in the intestine, we quantified *Cb* CFU in fecal samples from infected mice. Goup I *Cb* produces lipase, which enables the identification of *Cb* colonies by formation of a pearly layer on egg yolk Brucella HK agar (Fig. 2a). Colony PCR targeting the type A BoNT gene (*bont/A*) confirmed that all tested lipase-positive colonies were *Cb,* whereas lipase-negative colonies were negative for *Cb* (Fig. 2b). These results indicate that the lipase-positive colonies detected in fecal samples represented *Cb*. In AV-treated mice, lipase-positive colonies were detected as early as day 1 after spore administration. The number of colonies increased over time, reaching approximately 1 × 10 □ CFU/g feces on day 3 and 1 × 10 □ CFU/g feces on day 5 (Fig. 2c). In contrast, only a slight increase in CFU was observed in control mice on day 1, and lipase-positive colonies were undetectable thereafter. These results indicate *Cb* rapidly shed and did not persist in the control mice. *Cb* growth in AV-treated mice was also confirmed by quantitative PCR detection of the *bont/A* gene in fecal extracts (Fig. S2). To determine whether *Cb* existed primarily as spores or vegetative cells in the intestine, fecal samples were heat-treated before culture. Lipase-positive colonies were not detected in heat-treated fecal samples, suggesting that *Cb* was present mainly as vegetative cells, rather than heat-resistant spores, in the intestine of AV-treated mice on days 3 and 5 (Fig. 2d and Fig. S3). In contrast, at 10 h after spore administration, approximately 1 × 10□CFU/g feces of lipase-positive colonies were detected in both control and AV-treated mice, and colonies remained detectable after heat treatment (Fig.S4a). These results suggest that, at early time points after administration, many of the detected *Cb* were spores that had passed through the intestine without germination. Together, these results indicate that AV-induced dysbiosis permits sustained intestinal colonization and proliferation of *Cb*.

**Figure 2.**
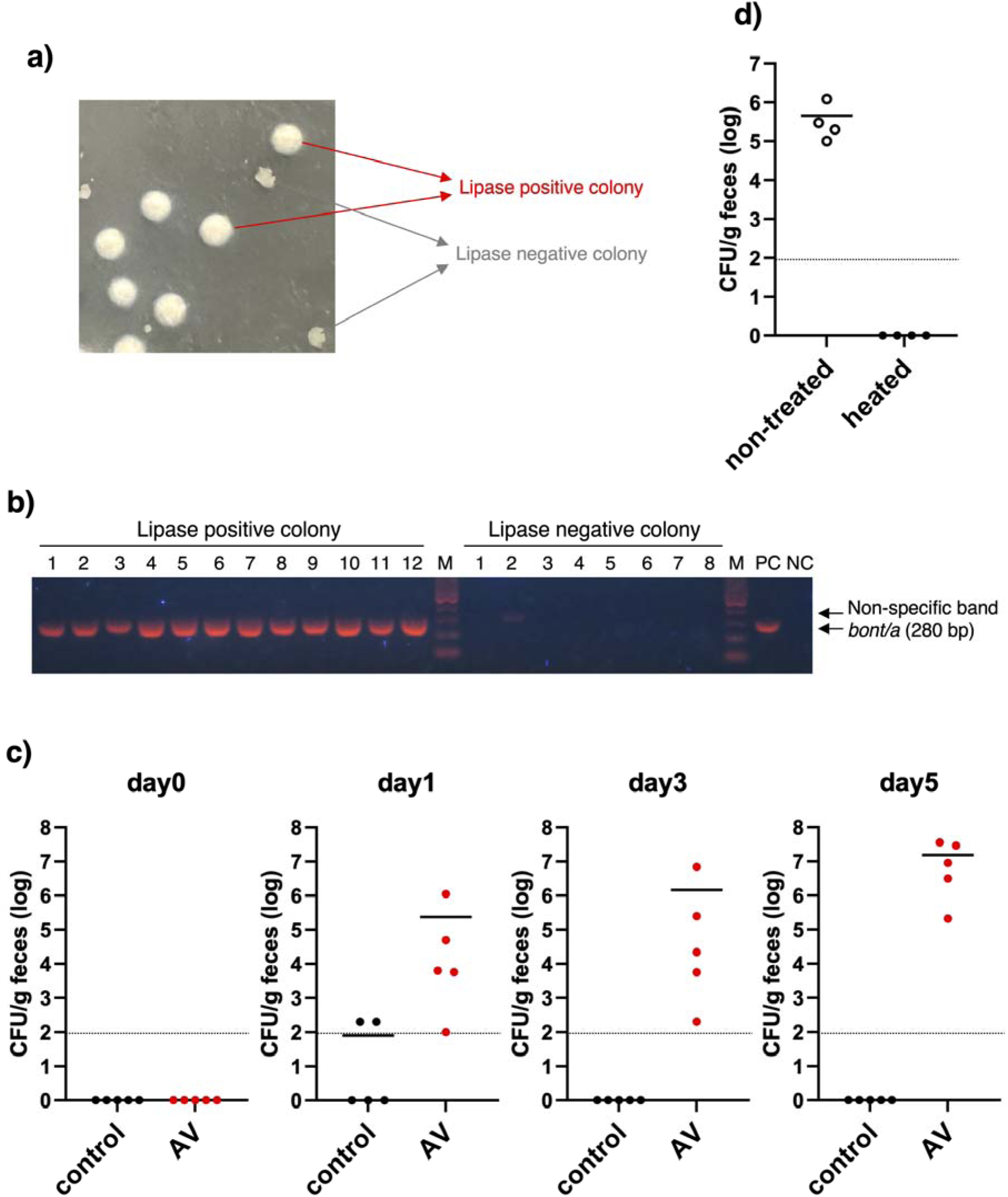
Detection of *Cb* cells in the fecal samples. **a** Representative image of lipase-positive colonies. Prepared fecal samples were cultured on egg-yolk containing Brucella HK agar. The plates were incubated anaerobically for 2 days at 37°C. Lipase-positive colonies were identified by the presence of a characteristic pearly layer. **b** Confirmation of the *bont/A* gene in lipase-positive colonies by colony PCR. Lipase-positive or lipase-negative colonies were picked and analyzed by colony PCR using specific primers for *bont/A* gene. M, molecular marker, PC, positive control (DNA of strain 62A), NC, negative control (water). **c** Quantification of lipase-positive colonies in fecal samples collected from control and AV-treated mice (*n* = 5) on days 0, 1, 3, and 5 after spore administration. Prepared fecal samples were seeded on the Brucella HK agar with egg-yolk. The plates were incubated anaerobically for 2 days at 37°C, and then lipase positive colonies were counted. **d** Prepared fecal samples on day 3 or 5 (2 samples each) were heated (80°C, 20 min) and then seeded on the Brucella HK agar with egg-yolk. The plates were incubated anaerobically for 2 days at 37°C, and then lipase positive colonies were counted. **cd** Each dot represents an individual mouse. The dotted line indicates the limit of detection. Data are representative of two independent experiments.

### *Cb* actively produces BoNT during infection in AV-treated mice

We next quantified the amount of serotype A BoNT (BoNT/A) in fecal samples using a BoNT/A-specific sandwich ELISA (Fig. S5). A significant increase in BoNT/A levels was observed in fecal samples from AV-treated mice on day 5 (Fig. 3a), indicating that large amounts of BoNT/A accumulated following expansion of the intestinal *Cb* population. In contrast, BoNT/A was not clearly detected in fecal samples from control mice, including on day 1, when *Cb* colonies were detected. To test the biological activity of BoNT/A in the fecal samples, we injected fecal supernatants intraperitoneally into mice. Fecal supernatants from AV-treated mice on day 3 showed toxic activity of ≥4,000 lethal dose 50% (LD50)/g, whereas those from control mice showed no detectable toxic activity, even at the lowest dilution tested, a 10-fold dilution (Fig. 3b). Together, these results demonstrate that AV-induced *Cb* colonization and expansion lead to the production of large amounts of biologically active BoNT/A in the intestine and subsequent disease progression.

**Figure 3.**
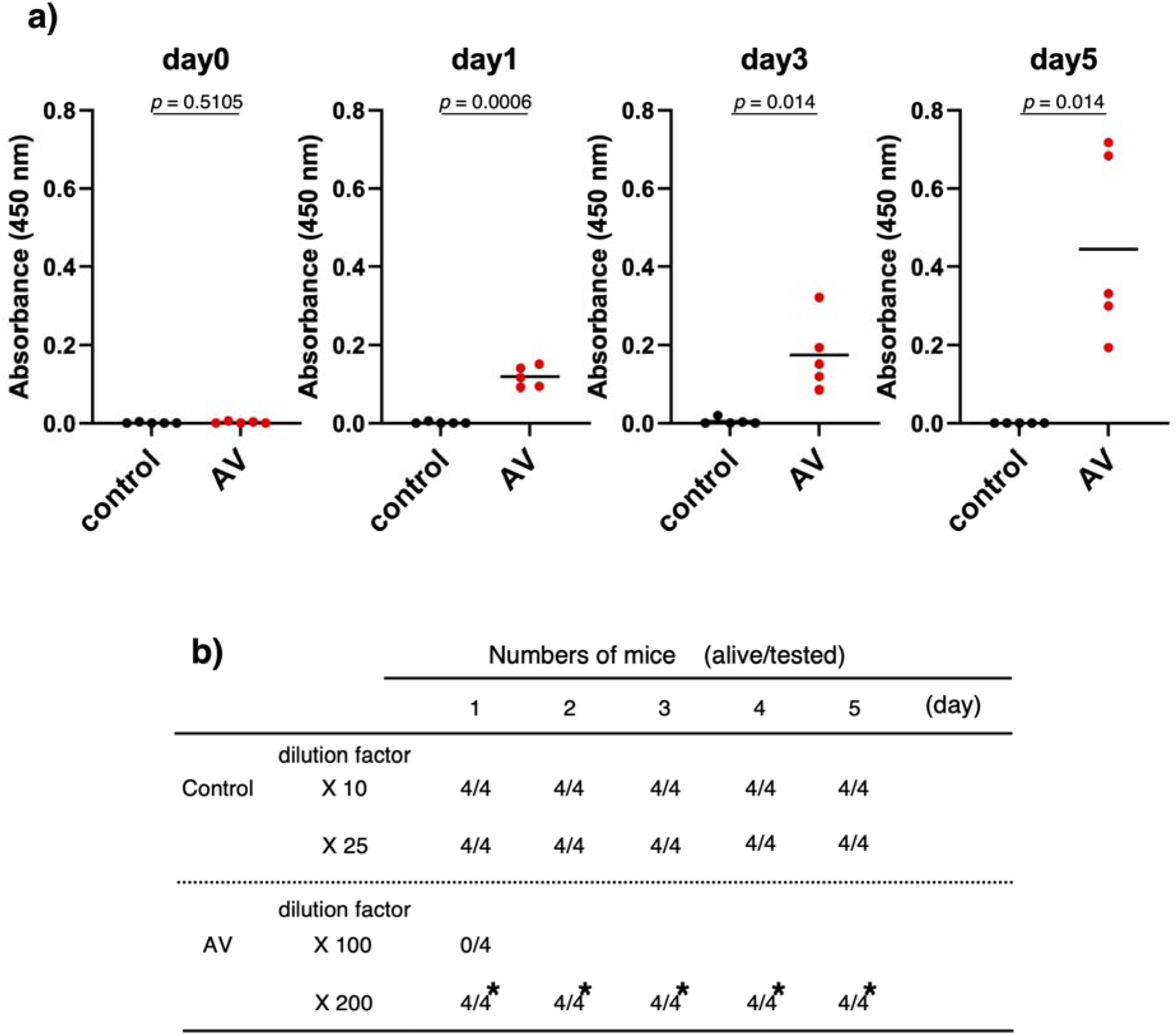
Detection of BoNT/A in the fecal samples. a BoNT/A levels in fecal samples were measured by sandwich ELISA. Supernatants of fecal samples (*n* = 5) were added to plates coated with mouse anti-BoNT/A monoclonal antibody. After washing, bound BoNT/A was detected by rabbit anti-BoNT/A polyclonal antibody and HRP-conjugated anti-rabbit IgG antibody. Each dot represents an individual mouse. The data were analyzed by two-tailed Welch’s t-test. **b** Toxic activity of fecal supernatants collected on day 3 was evaluated by mouse bioassay. Diluted fecal supernatants were intraperitoneally injected into mice (*n* = 4). Mice were monitored for morbidity and mortality for 5 days. Asterisks indicate samples for which all mice developed botulism symptoms. Data are representative of two independent experiments.

### Identification of the intestinal sites of *Cb* colonization and BoNT/A production

To identify the intestinal sites where *Cb* grows and BoNT/A accumulates, we quantified *Cb* CFU and BoNT/A levels in the small intestine, cecum, and colon of AV-treated mice on day 4 after spore administration. All mice showed mild or moderate symptoms on day 4. *Cb* was detected at approximately 1 × 10 □ CFU/g in the cecum and colon, whereas it was undetectable in the small intestine (Fig. 4a). Consistent with the distribution of *Cb* colonization, BoNT/A was detected in the cecum and colon (Fig. 4b). These intestinal contents showed toxic activity of ≥4,000 LD50/g, indicating the presence of biologically active BoNT/A (Fig. 4c). In contrast, only low levels of BoNT/A and weak toxic activity were detected in the small intestine. Furthermore, analysis of intestinal contents from mice immediately after disease onset (mild symptoms) and shortly before death (severe symptoms) showed similar patterns of toxic activity in the small intestine, cecum, and colon (Fig. S6). These results indicate that *Cb* primarily colonizes and expands in the cecum and colon of AV-treated mice, leading to the accumulation of biologically active BoNT/A at these sites.

**Figure 4.**
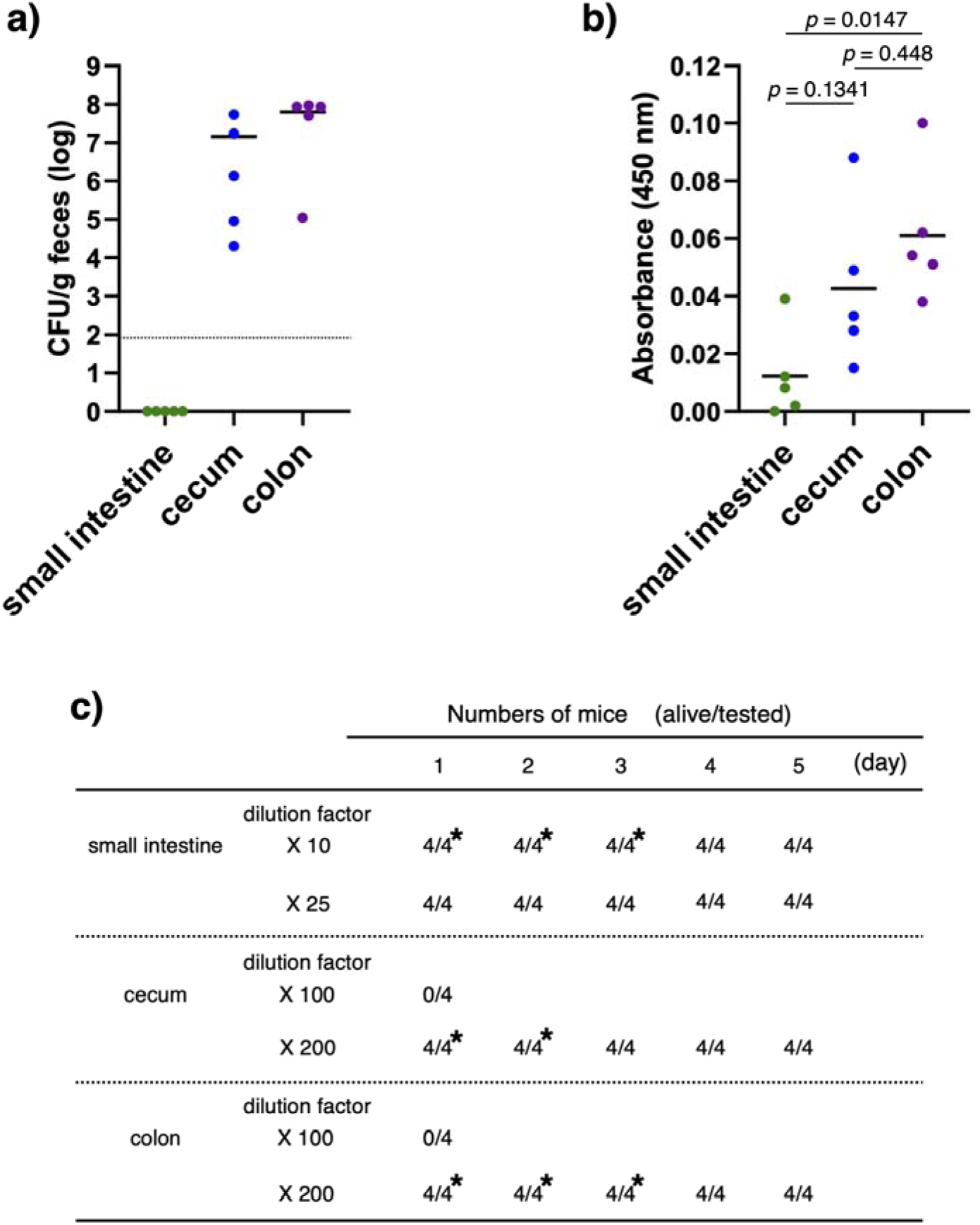
Tissue distribution of *Cb* cells and BoNT/A in infected mice. **abc** Spores (1 × 10^6^ CFU) were intragastrically administered to AV-treated mice (*n* = 5), and intestinal contents were collected on day 4. **a** Quantification of lipase-positive colonies in intestinal contents from the small intestine, cecum, and colon. Diluted intestinal contents were seeded on the Brucella HK agar with egg-yolk. The plates were incubated anaerobically for 2 days at 37°C, and then lipase-positive colonies were counted. Each dot represents an individual mouse. The dotted line indicates the limit of detection. **b** BoNT/A levels in intestinal contents were measured by sandwich ELISA using anti-BoNT/A antibodies. Intestinal contents supernatants were added to plates coated with mouse anti-BoNT/A monoclonal antibody. After washing, bound BoNT/A was detected by rabbit anti-BoNT/A polyclonal antibody and HRP-conjugated anti-rabbit IgG antibody. Each dot represents an individual mouse. The data were analyzed by one-way ANOVA followed by Tukey’s multiple-comparison test. **c** Toxic activity of intestinal content supernatants was evaluated by mouse bioassay. Diluted intestinal content supernatants were intraperitoneally injected into mice (*n* = 4). Mice were monitored for morbidity and mortality for 5 days. Asterisks indicate samples for which all mice developed botulism symptoms. Data are representative of two independent experiments.

### Application of the intestinal botulism mouse model to multiple *Cb* strains

*Cb* 62A is a long-standing laboratory strain producing BoNT/A1 (30). We also tested three additional *Cb* strains, including clinical strains isolated from infant botulism, to confirm the broad applicability of this mouse model: strains 7I03-H, a serotype A2 strain isolated from infant botulism (31); Okra, a serotype B1 strain isolated from foodborne botulism (32); and Osaka05, a serotype B6 strain isolated from infant botulism (33). These strains, including 62A, belong to Group I *Cb*, the group most commonly associated with infant botulism. All tested strains colonized in AV-treated mice and developed botulism symptoms (Fig. 5a–c). The bacterial abundance of *Cb* in fecal samples were similar among the strains at approximately 1 × 10 □ to 1 × 10 □ CFU/g on day 3 or day 5 (Fig. 2c and 5). The presence of the corresponding *bont* genes in lipase-positive colonies was confirmed by serotype-specific PCR (Fig. S7). Notably, *Cb* Okra showed the fastest onset, reaching a disease score of 2 by day 3–4 compared with 62A (Day 4), 7I03-H (Day 6), and Osaka05 (Day 6) (Fig. 1d and 5). These results demonstrate that this mouse model is broadly applicable to the analysis of intestinal botulism caused by multiple Group I *Cb* strains with different serotypes and clinical origins.

**Figure 5.**
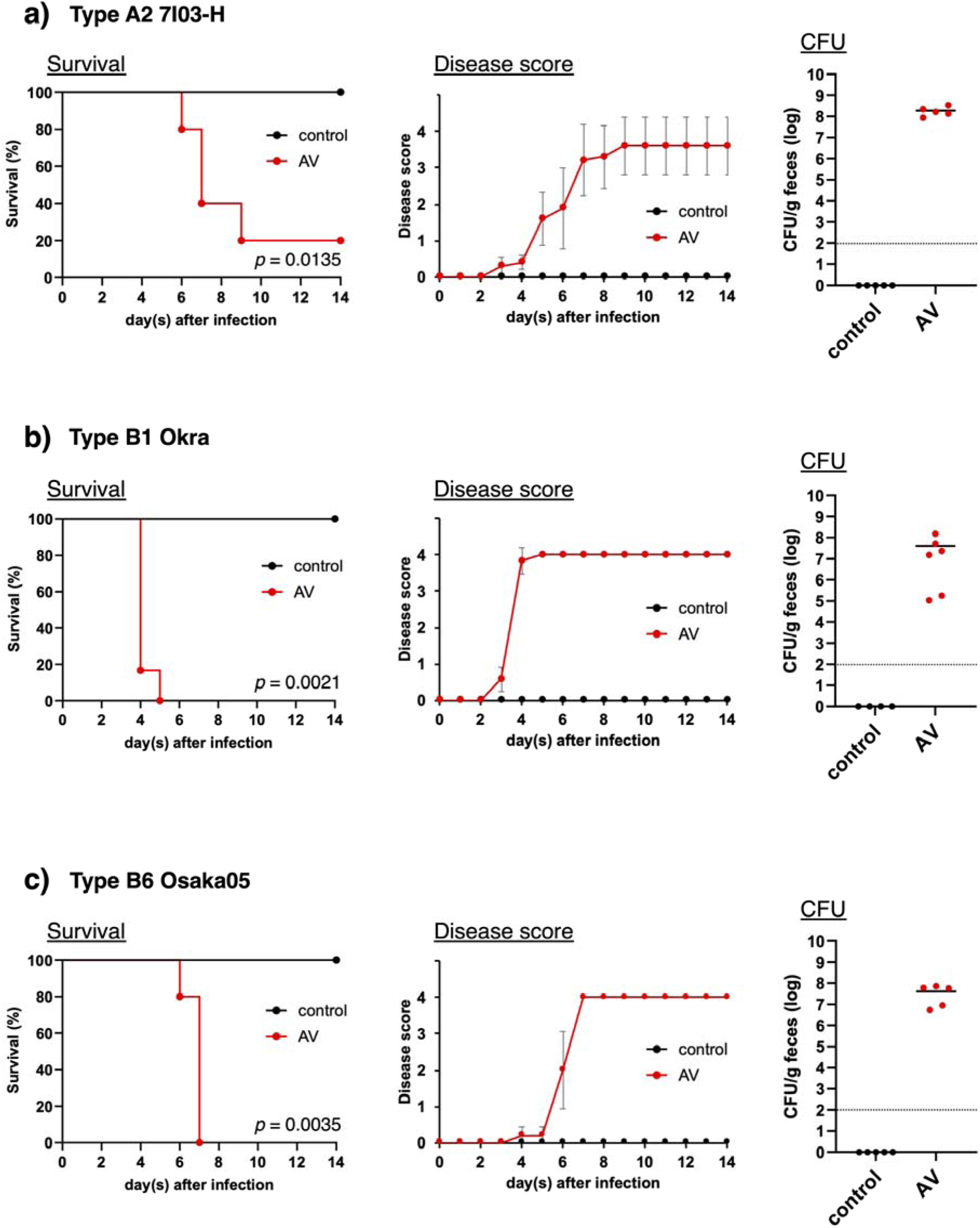
Multiple *Cb* strains can cause intestinal botulism in AV-treated mice. **a-c** AV-treated or control mice were intragastrically administered spores (1 × 10^6^ CFU) of strain 7I03-H, type A2 (**a**) (*n* = 5); Okra, type B1 (**b**) (control *n* = 4, AV *n* = 6); or Osaka05, type B6 (**c**) (*n* = 5). Spore-administered mice were monitored for survival and disease scores during a 2-week observation period. Statistical analysis of survival was performed using the log-rank test. For disease scores, data are presented as mean ± s.d. Fecal samples were collected on day 3 for Okra–administered mice and on day 5 for 7I03-H– and Osaka05–administered mice. Lipase-positive colonies in fecal samples were counted. Each dot represents an individual mouse. The dotted line indicates the limit of detection.

## Discussion

The intestinal microbiota is closely associated with susceptibility to intestinal infection by pathogenic bacteria (29,34–39). In the case of *Cb*, previous studies have shown that neonatal mice, which have immature intestinal microbiota, and germ-free mice are susceptible to infection following oral administration of botulinum spores (13–16). In addition, it is reported that *Cb* colonization and the development of intestinal botulism occurred in mice treated with a combination of erythromycin and kanamycin or metronidazole (17,18).

In the present study, we established an antibiotic-treated mouse model of intestinal botulism. In contrast to previous reports, we did not observe clear botulism symptoms in mice treated with metronidazole or with the combination of erythromycin and kanamycin (Fig. S1a). Because the previous studies were conducted nearly half a century ago, differences in animal housing conditions, microbiological status, and experimental environments over time may have contributed to the discrepancy between their findings and ours. This discrepancy may also be explained by differences in experimental conditions, including antibiotic administration conditions, mouse strain, age, housing environment, diet, and the baseline composition of the intestinal microbiota. In particular, intestinal microbiota can vary substantially among animal facilities, and such differences may influence both the effect of antibiotics on microbial communities and the efficiency of *Cb* colonization. In contrast, treatment with the combination of ampicillin and vancomycin rendered mice susceptible to intestinal botulism following intragastric administration of *Cb* spores (Fig. S1, 1cd). These findings suggest that AV treatment creates an intestinal environment that permits *Cb* colonization and disease development, providing a reproducible model for analyzing the pathogenesis of intestinal botulism. Ampicillin is a β-lactam antibiotic active against both Gram-positive and Gram-negative bacteria, whereas vancomycin is a glycopeptide antibiotic mainly active against Gram-positive bacteria, including antimicrobial-resistant bacteria. Previous studies have reported that AV treatment efficiently disrupts the intestinal microbiota. Indeed, AV-treated mice develop intestinal dysbiosis and show increased susceptibility to viral infection (40). These findings suggest that AV treatment reduces total bacterial abundance, markedly limits microbial diversity, and thereby increases susceptibility to intestinal pathogens. In this study, AV treatment markedly reduced total bacterial abundance in the mouse intestine (Fig. 1b) and created an environment permissive for *Cb* colonization and intestinal botulism. This may be due to depletion of bacterial populations that normally provide colonization resistance against *Cb*. AV-induced dysbiosis may also reduce microbial competition or alter the intestinal metabolic environment, thereby facilitating *Cb* persistence and expansion. The detailed relationship between *Cb* and the intestinal microbiota remains unclear. However, because susceptibility to intestinal botulism varied depending on the antibiotics used, comparative microbiota analysis after different antibiotic treatments may help identify bacterial taxa or microbial functions that protect against *Cb* colonization. This approach may contribute to the development of microbiota-based strategies for preventing intestinal botulism.

Control mice showed no botulism symptoms throughout the observation period, although a slight increase in CFU was observed in fecal samples at 10 h and on day 1 after spore administration (Fig. 2c and Fig. S4a). At 10 h after spore administration, fecal samples from control and AV-treated mice showed comparable CFU levels, which did not markedly decrease after heat treatment. Furthermore, no toxic activity was detected in these samples (Fig. S4b). In both groups, this transient increase likely reflects the passage of intragastrically administered spores through the intestine rather than successful colonization. In control mice, the absence of detectable *Cb* in fecal samples after day 3 suggests that mature intestinal microbiota may prevent *Cb* colonization, leading to elimination of the administered spores from the intestinal tract. In contrast, AV-treated mice showed a time-dependent increase in CFU and BoNT/A levels in fecal samples, suggesting that intragastrically administered spores germinated and expanded in the intestine under dysbiotic conditions. Consistent with this interpretation, *Cb* was present mainly as heat-sensitive vegetative cells rather than heat-resistant spores in the intestine of AV-treated mice (Fig. 2d and Fig. S3). BoNT/A levels increased after expansion of the intestinal *Cb* population, particularly on days 3–5, before the appearance of obvious symptoms (Fig. 3a). These findings suggest that disease onset requires not only intestinal colonization by *Cb* but also accumulation of sufficient amounts of biologically active BoNT/A. Supporting this idea, intraperitoneal administration of anti-BoNT/A antibody inhibited the onset botulism in this model (data not shown). The maximum CFU level detected in feces was 1.5 × 10 □ CFU/g on day 5, whereas toxic activity reached 4,000–8,000 LD50/g on day 3 (Fig. 2c and Fig. 3b). Toxic activity in fecal samples on day 5 could not be systematically analyzed because only small amounts of feces could be collected from severely affected AV-treated mice. However, an analyzable day 5 sample from one mouse showed toxic activity of 16,000–32,000 LD50/g, suggesting that expansion of the intestinal *Cb* population leads to accumulation of large amounts of biologically active BoNT/A, which likely contributes to disease progression. These results demonstrate that AV-induced dysbiosis permits intestinal expansion of *Cb*, leading to accumulation of biologically active BoNT/A before the onset of clinical symptoms. This temporal analysis distinguishes our model from previous susceptibility-based studies and provides a useful platform for investigating the pathogenesis of intestinal botulism. Moreover, the CFU and toxic activity detected in fecal samples from our mouse model were comparable to those previously reported in fecal samples from patients with adult intestinal toxemia botulism (41–43). These findings suggest that our model partially recapitulates key pathological features of human intestinal botulism, including intestinal *Cb* expansion and accumulation of biologically active BoNT/A. Real-time PCR (qPCR) using *bont/A*-specific primers also detected *Cb* in fecal samples, and the results were consistent with culture-based CFU analysis (Fig. S2). Unlike CFU analysis, qPCR cannot determine bacterial viability. However, qPCR is simple, highly specific, and quantitative, and may therefore serve as a useful complementary method for monitoring intestinal *Cb* burden. In combination with toxin detection, fecal qPCR may help evaluate the progression of intestinal botulism and support diagnostic assessment.

*Cb* growth was observed mainly in the cecum and colon, but not in the small intestine (Fig. 4a). In contrast, a small amount of BoNT/A and weak toxic activity, less than 400 LD50/g, were detected in the intestinal contents of the small intestine from immediately after disease onset to shortly before death (Fig. 4b,c, S6), consistent with a previous report (18). There are several possible explanations for this observation. First, BoNT/A produced in the cecum and colon may move back into the small intestine because of dysfunction or paralysis of the ileocecal valve during disease progression. Second, a small number of *Cb* cells below the detection limit of the culture assay may produce BoNT/A in the small intestine. Previous studies have suggested that BoNT is absorbed more efficiently from the small intestine than from the large intestine (44–46). In foodborne botulism, BoNT/A has been reported to exploit microfold cells in the small intestine to enter the host and exert neurotoxicity (47). Therefore, even a small amount of BoNT/A detected in the small intestine may contribute to the development of systemic botulism symptoms. These findings suggest that the cecum and colon are the major sites of *Cb* expansion and BoNT/A accumulation in this model, whereas the small intestine may play an important role as a site of BoNT/A absorption. Further studies are needed to clarify the mechanism of BoNT/A transport and absorption in the intestinal botulism mouse model.

Human intestinal botulism is mainly caused by serotypes A and B. In this study, in addition to the type A1 strain 62A, we used two strains isolated from infant botulism, type A2 strain 7I03-H and type B6 strain Osaka05, and one type B1 strain isolated from foodborne botulism, Okra, to evaluate the versatility of this intestinal botulism mouse model. These strains possess different BoNT gene clusters, namely the *ha* or *orfX* cluster (48–50), whose associated proteins are known to contribute to the oral toxicity of BoNT complexes (47,51–56). All AV-treated mice administered spores of each strain developed botulism symptoms (Fig. 5a-c), indicating that this model is applicable to intestinal botulism caused by multiple *Cb* strains with different serotypes, subtypes, clinical origins, and toxin gene cluster types. Among the tested strains, AV-treated mice administered type B1 strain Okra showed shorter survival times and a shorter interval from symptom onset to death than mice administered the other strains (Fig. 5b). Previous studies have reported that the toxin complex produced by type B1 Okra has relatively high oral toxicity (4,57,58). Therefore, the more rapid disease progression observed in Okra-administered mice may reflect strain-dependent differences in the oral toxicity of BoNT complexes. These findings suggest that the properties of toxin complexes may contribute not only to the severity of foodborne botulism but also to the progression of intestinal botulism. Thus, this mouse model can be used to compare the pathogenicity of diverse *Cb* strains and to evaluate how differences in BoNT complexes and toxin-associated proteins influence intestinal botulism.

Taken together, our findings establish a reproducible antibiotic-treated mouse model that enables quantitative analysis of the entire pathological course of intestinal botulism, from *Cb* colonization and intestinal expansion to BoNT accumulation, symptom onset, and death. This model is applicable to multiple *Cb* strains with different serotypes and clinical origins, allowing comparative analysis of strain-dependent pathogenicity. Importantly, using this model, we clarified how intestinal bacterial burden, biologically active toxin accumulation, and progression of botulism symptoms are temporally linked during disease development. These findings advance our understanding of the mechanisms underlying intestinal botulism pathogenesis and may contribute to the development of novel preventive and therapeutic strategies.

## Materials and methods

### Biosafety

Our laboratory at Kanazawa University is authorized by the Minister of Health, Labour and Welfare of Japan to possess and handle *Clostridium botulinum* and botulinum neurotoxins, which are classified as Class II pathogens under the relevant Cabinet Order of Japan. The research program and all personnel involved in this study were also approved and monitored by the Kanazawa University Biosafety Committee for Pathogens.

### Ethics statement

Animal studies were conducted under the applicable laws and guidelines for the care and use of laboratory animals at the Kanazawa University. They were approved by the Animal

Experiment Committee of the Kanazawa University (AP-214246 and AP-214252).

### Preparation of botulinum spores

*C*. *botulinum* strain 62A, 7I03-H, Okra, and Osaka05 were used in this study. *Cb* was cultured in Tryptone-peptone-glucose-yeast (TPGY) medium (BD Difco, Franklin Lakes, NJ): 5% Trypticase peptone (BD Difco), 0.5% Bacto peptone (BD Difco), 0.4% glucose (Fujifilm, Osaka, Japan), 2% yeast extract (Sigma-Aldrich, St Louis, MO), 0.1% L-cysteine HCl (Fujifilm) (pH 7.3 −7.4) overnight at 37°C in an anaerobic chamber (82% nitrogen, 10% hydrogen, 8% carbon dioxide; ANAEROBOX, HIRASAWA, Tokyo, Japan). A 0.1 ml aliquot of cultures (diluted 1:10^4^–1:10^6^) were seeded on TPGY plates and cultured for 10–14 days at 37°C under anaerobic conditions. The colonies were collected from TPGY plates in ice-cold H_2_O and centrifuged (12,000 xg, 20 min, 4°C). The spore pellets were washed with sterilized ice-cold H_2_O by repeated centrifugation at 4°C for 10 min, with the centrifugation speed gradually reduced in the following order: 8,000 xg, 6,000 xg, 4,000 xg, 3,000 xg, and 2,000 xg, and then resuspended in 20% solution of Gastrografin (Bayer AG, Leverkusen, Germany) and layered over 50% Gastrografin solution to create a density gradient (59). The spore suspension was centrifuged (10,000 xg, 30 min, 4°C) and the layer of debris and extra Gastrografin solution were removed. The spore pellets were then washed with ice-cold H_2_O and then resuspended in sterilized H_2_O. The spore formation rate and purity were measured by phase contrast microscopy. We confirmed that more than 90% of *Cb* were present as spores.

### Antibiotics treatment

Metronidazole (Fujifilm, 1.0 mg/ml), a combination of erythromycin (Fujifilm, 1.0 mg/ml) and kanamycin (Sigma-Aldrich, 1.0 mg/ml), or combination of ampicillin (Sigma-Aldrich, 1.0 mg/ml) and vancomycin (Fujifilm, 0.5 mg/ml) were prepared in water, respectively. To make these solutions easy to drink, sucrose (Fujifilm, 10 mg/ml) was added to each solution. These antibiotic solution or sucrose solution (control) were administered via drinking water to BALB/c adult mice (female, 7–8 weeks, SLC, Hamamatsu, Japan) for 1 week, and then replaced with sterilized water to allow antibiotic wash out in mice for 36–48 h before administration of spores. In the case of metronidazole, 0.5 mg was administered by oral gavage once a day for 7 days. In this study, the combination of ampicillin and vancomycin was mainly used for dysbiosis.

### Spore challenge and disease scoring

*Cb* spore suspension was heated (80°C, 20 min) and diluted to 1 × 10^7^ CFU/ml with sterilized water. A 0.1 ml aliquot of spore suspension (1 × 10^6^ CFU) was intragastrically administered to antibiotic-treated or control mice. After administration, mice were housed individually in mesh-bottom cages placed in an individually ventilated cage rack to minimize coprophagy. Mice were monitored for morbidity and mortality during a 2-weeks observation period. Disease severity was scored for 2 weeks with symptoms ranging from “score: 0, no symptoms” to scores of “1: mild”, “2: moderate”, “3: severe”, and “4: death” defined by an increasing extent of botulism symptoms (ruffled fur, muscle weakness, limb weakness, and respiratory failure).

### Determination of CFU

At each time point, approximately 200 mg of feces was collected from each mouse, and a portion of each sample was randomly selected for subsequent analyses. For intestinal content samples, the small intestine, cecum, and colon were opened, and the contents were collected using a sterile spatula or other sterile instruments. The collected samples were weighed and then used for subsequent analyses. Fecal samples and intestinal contents were weighed and then suspended in sterile phosphate-buffered saline (PBS) to 0.1 g/ml. For heat-treated samples, suspensions were heated at 80°C for 20 min before dilution. Each suspension was further diluted with PBS and 0.1 ml aliquots were plated on Brucella HK agar supplemented egg yolk. The plates were incubated anaerobically for 2 days at 37°C using the AnaeroPack system (Mitsubishi Gas Chemical Company, Inc., Tokyo, Japan), and then lipase-positive colonies, which showed a characteristic pearly layer, were counted. To confirm the presence of *Cb*, we performed the colony PCR using specific primers for *bont/A* (F 5’-TGCAGGACAAATGCAACCAGT-3’, R 5’-TCCACCCCAAAATGGTATTCC-3’) or *bont/B* (F 5’-CCT CCA TTT GCG AGA GGT ACG-3’, R 5’-CTC TTC GAG TGG AAC ACG TCT −3’) (60). CFU values were calculated and expressed as CFU per gram of fecal sample or intestinal content.

### Real-time PCR (quantitative PCR, qPCR)

Approximately 50 mg per fecal sample were collected into a 2-ml tube with 0.1-mm and 3.0-mm zirconia beads. The fecal samples were homogenized at 1,500 rpm for 10 min with a Shake Master Neo (Bio Medical Sciences, Japan) after adding Inhibit EX buffer from the QIAamp Fast DNA Stool Mini Kit (QIAGEN, Venlo, Nederlands). Genomic DNA was subsequently extracted with the kit according to the manufacturer’s protocol. Quantitative PCR was performed using a QuantStudio 3 real time PCR system (Thermo) with KOD SYBR qPCR Mix (TOYOBO). Total bacterial abundance was quantified using primers targeting the V3–V4 region of the bacterial 16S rRNA gene5’-CCTACGGGNGGCWGCAG-3’, 5’-GACTACHVGGGTATCTAATCC-3’, *Cb* burden in fecal samples from strain 62A-administered mice was quantified using *bont/A*-specific primers; 5’-GGGCCTAGAGGTAGCGTARTG-3’, 5’-TCTTYATTTCCAGAAGCATATTTT-3’ as described previously (61). Real-time PCR data were analyzed using the comparative Ct method. Relative values were calculated using the ΔΔCt method, with fecal samples from untreated control mice used as the reference.

Relative values were calculated as 1 / 2^(Ct_sample − Ct_reference), where Ct_sample represents the Ct value of each sample and Ct_reference represents the Ct value of fecal samples from untreated control mice.

### Toxin (BoNT/A) ELISA

BoNT/A levels were measured by sandwich ELISA. 96-well plates were coated with mouse anti-BoNT/A monoclonal antibody, kindly provided by Dr. Tomoko Kohda at Osaka Metropolitan University, (100 ng/well) in 50 mM carbonate buffer at pH 9.6 overnight at 4°C. Wells were washed three times with PBS containing 0.05% Tween20 (Sigma-Aldrich, St. Louis, MO, USA) (PBS-T) and blocked with 0.5% bovine serum albumin (BSASigma-Aldrich)/PBS-T for 1 h at 37°C. Supernatants of fecal samples or intestinal contents at 0.1 g/ml in PBS containing EDTA-free protease inhibitor (Roche, Basel, Switzerland), as described for the Mouse bioassay, were added to the wells and incubated for 1 h at 37°C. After washing with PBS-T, rabbit anti-BoNT/A polyclonal antibody generated in our laboratory (25 ng/well) was added and incubated for 1 h at 37°C. After another wash with PBS-T, horseradish peroxidase (HRP)-conjugated anti–rabbit IgG (Jackson ImmunoResearch, West Grove, PA, USA) was added and incubated for 1 h at 37°C. Plates were washed again with PBS-T and then incubated with TMB solution (Thermo Fisher Scientific, Waltham, MA, USA) for 20 min at 37°C. Stop solution (0.2 M sulfuric acid) was added, and absorbance at 450 nm was measured using MULTISKAN plate reader (Thermo Fisher Scientific).

### Mouse bioassay

Fecal samples and intestinal contents were prepared in sterile PBS containing EDTA-free protease inhibitor to 0.1 g/ml and centrifuged (6,000 xg, 20 min, room temperature). The supernatants were filtered (0.2 µm) and further diluted with 10 mM sodium phosphate buffer (pH 6.0) containing 0.1% gelatin. 0.5 ml of diluted samples were intraperitoneally injected into mice (ICR, 4–5 weeks, SLC). Mice were monitored for morbidity and mortality for 5 days. The toxic activity of each sample was estimated as LD50 per gram of feces or intestinal contents based on the dilution factor and mortality rate observed in the mouse bioassay.

### Statistical analysis

All statistical analyses were performed using GraphPad Prism software version 11.0.2. Statistical significance was evaluated using an unpaired Welch’s *t*-test, one-way ANOVA followed by Tukey’s multiple-comparison test, or the log-rank test, as appropriate. Differences were considered statistically significant at *p* < 0.05.

## Abbreviations

AV: ampicillin and vancomycin
BoNT: botulinum neurotoxin
BSA: bovine serum albumin
Cb: Clostridium botulinum
CFU: colony forming unit
EK: erythromycin and kanamycin
HRP: horseradish peroxidase
LD50: lethal dose 50%
Met: metronidazole
M-PTC: medium progenitor toxin complex
PBS: phosphate-buffered saline
PBS-T: PBS containing 0.05% Tween20
qPCR: quantitative PCR
BoNT/A: type A botulinum neurotoxin

## Author contributions

T.M. and Y.F. designed the study and analyzed the data. T.M. performed the most of the experiments and analyses. N.K., S.A., and K.S. optimized the conditions for the infection experiments. N.K. developed the qPCR assays and analyzed the data. A.Y. and K.I. contributed to data analysis. T.M. and Y.F. co-wrote the manuscript. All authors discussed the data and approved the final version of the manuscript.

## Acknowledgments

We thank S. Akagi, Y. Koino, H. Kuraoka, Y. Konoshita, and H. Honda (Department of Bacteriology, Graduate School of Medical Sciences, Kanazawa University) for their technical assistance and members of the Fujinaga laboratory for valuable discussions. We are grateful to K. Umeda (Division of Microbiology, Osaka Institute of Public Health) for providing strain Osaka05. We also thank Dr. T. Kohda (Osaka Metropolitan University) for kindly providing the anti-BoNT/A monoclonal antibody. During the preparation of this manuscript, the authors used ChatGPT, developed by OpenAI, for English language editing. This work was supported in part by Grants-in-Aid for Scientific Research from the Japan Society for the Promotion of Science KAKENHI Grant Numbers 21K07021, 21H02729, and 25K10359; the Japan Agency for Medical Research and Development, AMED, Grant Numbers 21fk0108101j0003, 22fk0108635h0401, and 25fk0108728h0001; and the Ohyama Health Foundation Inc.

