## Supplementary Material for "An Antibiotic-Treated Mouse Model Reveals the Progression of Intestinal Botulism"

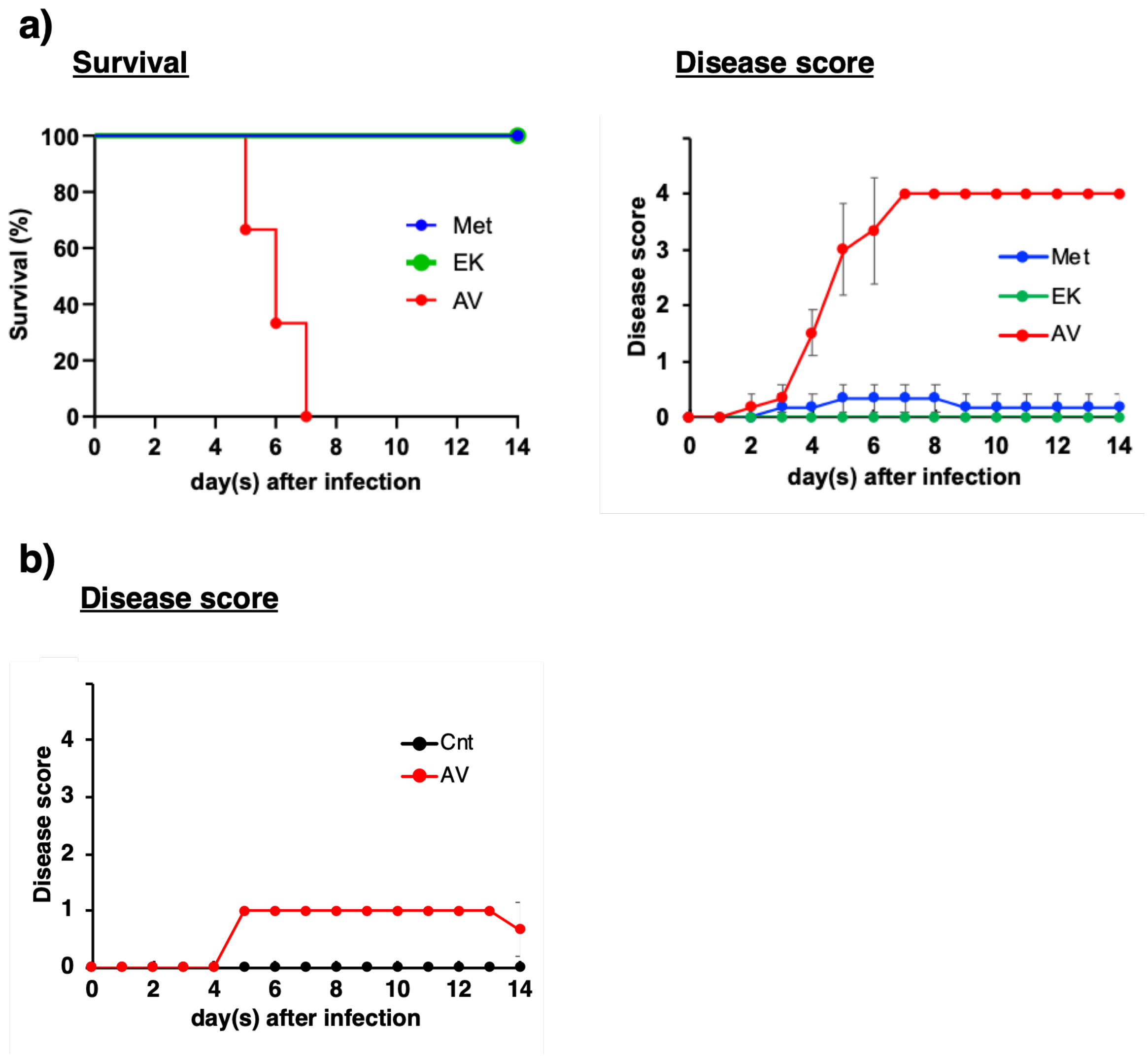


**Supplementary Figure 1. Comparative analysis of antibiotics for the intestinal botulism mouse model.**

**a** Mice were treated with metronidazole (Met), a combination of erythromycin and kanamycin (EK), or a combination of ampicillin and vancomycin (AV) before intragastric spore administration. After antibiotic washout, strain 62A spores (1 x $10^6$ CFU) were intragastrically administered to antibiotic-treated mice ($n$ = 3). Mice were monitored for morbidity, mortality, and survival during a 2-week observation period to determine which antibiotics reproducibly induced susceptibility to intestinal botulism. **b** AV-treated mice or control mice were orally administered strain 62A spores (1 × $10^5$ CFU) ($n$ = 3). Mice were monitored for 2 weeks after spore administration. All AV-treated mice developed botulism symptoms, although none died during the observation period. In contrast, control mice showed no symptoms throughout the observation period.

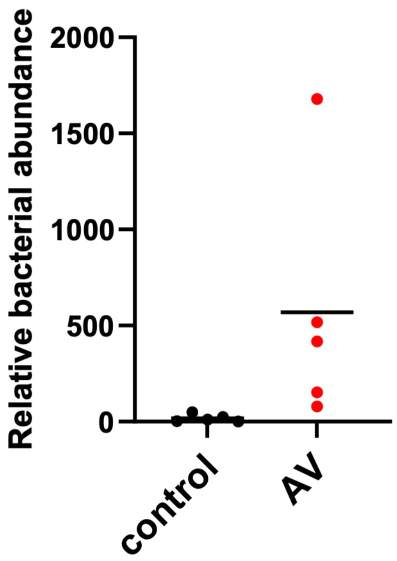


**Supplementary Figure 2. qPCR detection of *Cb* in fecal samples from infected mice.** DNA was extracted from fecal samples collected on day 3 (*n* = 5) using the QIAamp Fast DNA Stool Mini Kit. The *bont/A* gene was detected by qPCR using *bont/A*-specific primers to evaluate the intestinal burden of *Cb* in the mouse model. Each dot represents an individual mouse.

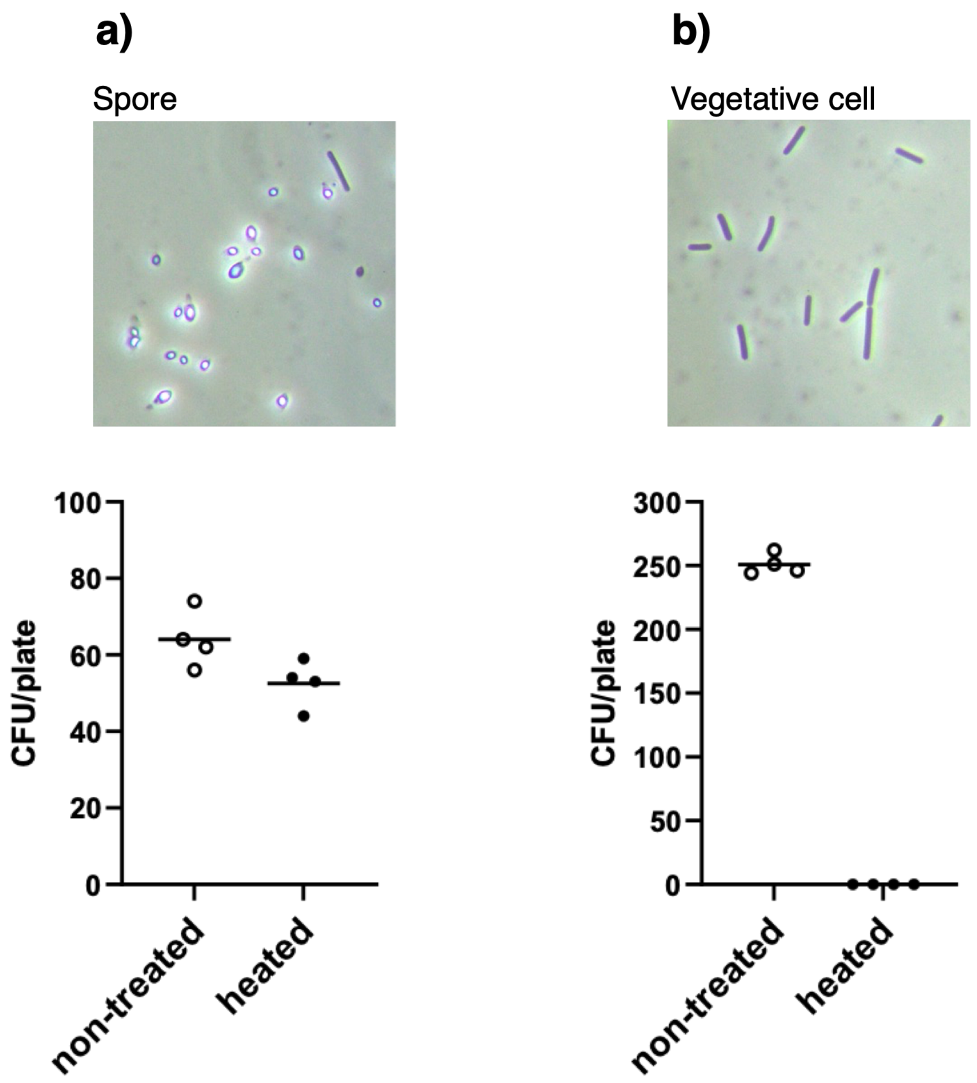


**Supplementary Figure 3. Heat resistance of *Cb* spores and heat sensitivity of vegetative cells.**

Spore suspensions (**a**) and vegetative cell suspensions (**b**) were heat-treated (80°C, 20 min) and analyzed by culture-based detection of lipase-positive colonies. The upper panels show phase-contrast microscopy images. Heat-treated spores retained colony-forming ability, whereas heat-treated vegetative cells did not form detectable colonies. These results support the interpretation that the heat-sensitive *Cb* cells detected in fecal samples from infected mice in Fig. 2d were predominantly vegetative cells rather than spores.

a)

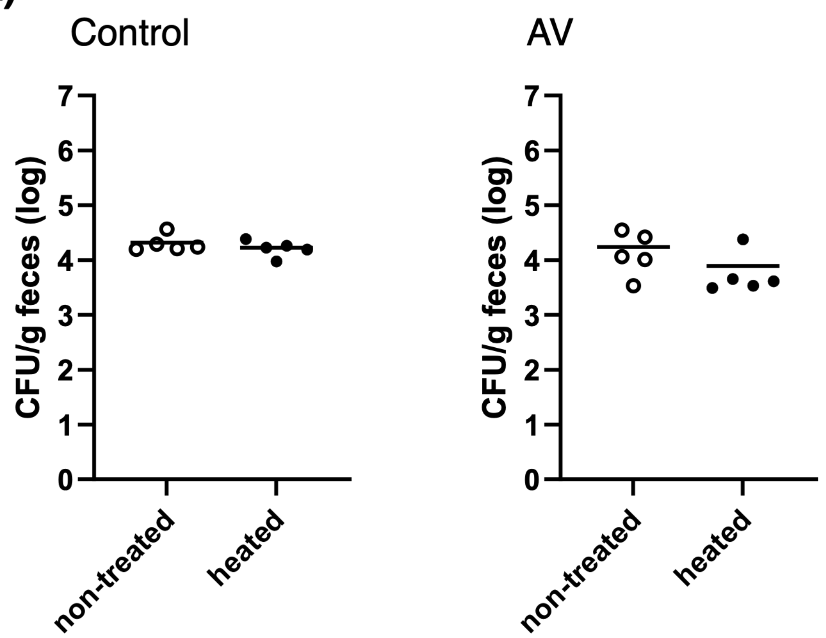


b)

| | | Numbers of mice (alive/tested) | | | | | |
|---|---|---|---|---|---|---|---|
| | | 1 | 2 | 3 | 4 | 5 | (day) |
| | dilution factor | | | | | | |
| Control | X 10 | 4/4 | 4/4 | 4/4 | 4/4 | 4/4 | |
| AV | X 10 | 4/4 | 4/4 | 4/4 | 4/4 | 4/4 | |

**Supplementary Figure 4. Analysis of the bacterial state of *Cb* and toxic activity in fecal samples collected 10 h after spore administration.**

**a** Fecal samples collected 10 h after spore administration were either heat-treated (80°C, 20 min) or left untreated and then seeded on the Brucella HK agar with egg-yolk. The plates were incubated anaerobically for 2 days at 37°C, and then lipase positive colonies were counted. Each dot represents an individual mouse. The dotted line indicates the limit of detection. **b** Toxic activity of fecal supernatants collected 10 h after spore administration was evaluated by mouse bioassay. Diluted fecal supernatants were intraperitoneally injected into mice ($n = 4$). Mice were monitored for morbidity and mortality for 5 days.

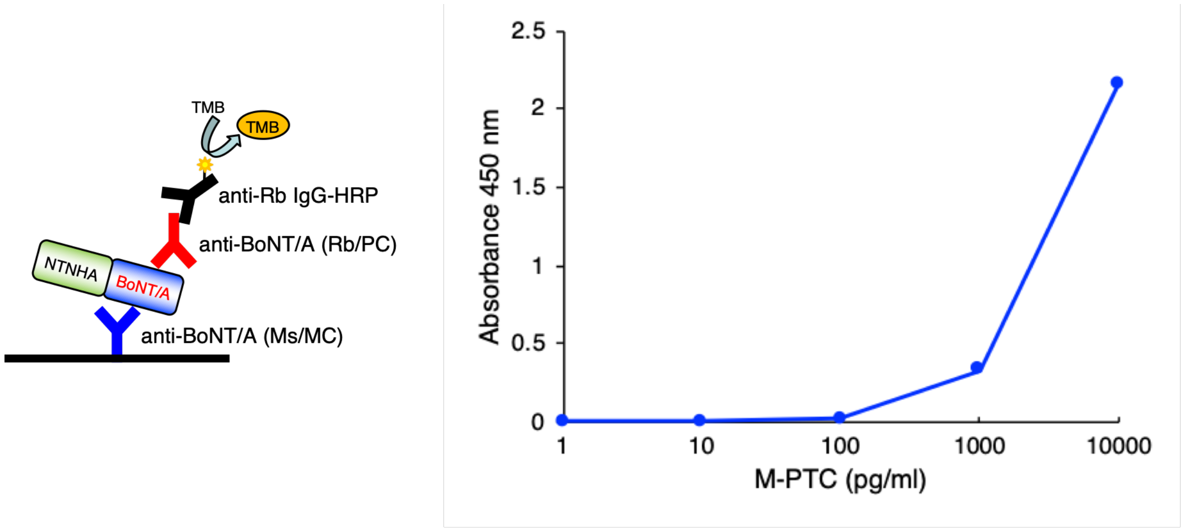


**Supplementary Figure 5. Detection of BoNT/A by sandwich ELISA.**

Detection sensitivity of the sandwich ELISA using purified serotype A medium progenitor toxin complex (M-PTC/A). M-PTC/A consists of BoNT/A and non-toxic non-hemagglutinin (NTNHA). BoNT/A in M-PTC/A was detected by sandwich ELISA using plates coated with a mouse anti-BoNT/A monoclonal antibody (Ms/MC). After blocking with BSA, M-PTC/A was added to the wells and incubated. Bound BoNT/A was detected using a rabbit anti-BoNT/A polyclonal antibody (Rb/PC), followed by an HRP-conjugated anti-rabbit IgG antibody. The signal was developed with TMB substrate solution, stopped with stop solution, and measured at 450 nm. Detection of purified M-PTC/A confirmed that this ELISA system can detect BoNT/A-containing toxin complexes. Data are presented as the mean of duplicate wells and representative of at least two independent experiments.

**Disease score 1.0 (mild)**

| | | Numbers of mice (alive/tested) | | | | | |
|---|---|---|---|---|---|---|---|
| | | 1 | 2 | 3 | 4 | 5 | (day) |
| small intestine | dilution factor X 10 | 4/4* | 4/4* | 4/4* | 4/4* | 4/4* | |
| cecum | dilution factor X 100 | 1/4* | 0/4 | | | | |
| colon | dilution factor X 100 | 3/4* | 1/4* | 0/4 | | | |

**Disease score 3.0 (severe)**

| | | Numbers of mice (alive/tested) | | | | | |
|---|---|---|---|---|---|---|---|
| | | 1 | 2 | 3 | 4 | 5 | (day) |
| small intestine | dilution factor X 10 | 4/4* | 4/4* | 4/4* | 4/4* | 4/4 | |
| cecum | dilution factor X 100 | 0/4 | | | | | |
| colon | dilution factor X 100 | 4/4* | 1/4* | 0/4 | | | |

**Supplementary Figure 6. Analysis of toxic activity in intestinal contents.**

Toxic activity in intestinal content supernatants from mice with mild symptoms (disease score 1) or severe symptoms (disease score 3) was evaluated by mouse bioassay. Diluted intestinal content supernatants were intraperitoneally injected into mice ($n = 4$). Mice were monitored for morbidity and mortality for 5 days. Asterisks indicate samples for which all mice developed botulism symptoms.

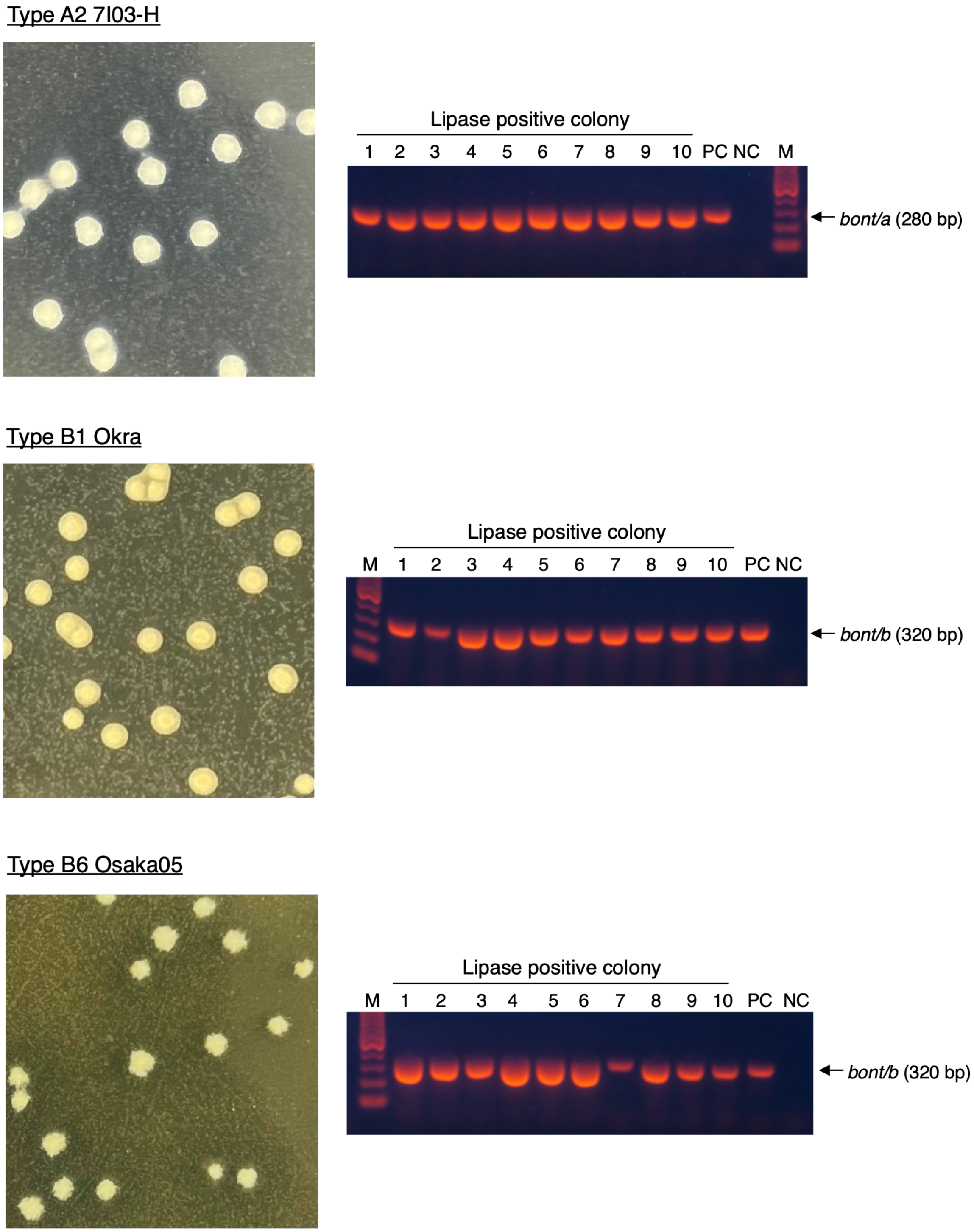


**Supplementary Figure 7. Confirmation of *bont* genes in lipase-positive colonies isolated**

**from fecal samples of mice infected with strains 7I03-H, Okra, or Osaka05.**

Lipase-positive colonies isolated from fecal samples of infected mice were analyzed by colony PCR using *bont*-specific primers. All tested lipase-positive colonies carried the corresponding *bont* gene, confirming that these colonies were *Cb*. The expected PCR product sizes were 280 bp for strain 7I03-H and 320 bp for strains Okra and Osaka05. M, molecular marker; PC, positive control using DNA from *Cb*; NC, negative control using water.
